# SGCI (Smartwatch Gait Coordination Index): New Measure for Human Gait Utilizing Smartwatch Sensor

**DOI:** 10.1101/2022.12.19.521019

**Authors:** Sumin Han, Robert Paul

**Affiliations:** Korea International School, Pangyo, South Korea; HS Math & AP Calculus BC Department Head in Korea International School, Pangyo, South Korea

## Abstract

Human walking reflects the state of human health. Numerous medical studies have been conducted to analyze walking patterns and to diagnose disease progression. However, this process requires expensive equipment and considerable time and manpower. Smartwatches are equipped with gyro sensors to detect human movements and graph-walking patterns. To measure the abnormality in walking using this graph, we developed a smartwatch gait coordination index (SGCI) and examined its usefulness. The phase coordination index was applied to analyze arm movements. Based on previous studies, the PCI formula was applied to graphs obtained from arm movements, showing that arm and leg movements during walking are correlated with each other. To prove this, a smartwatch was worn on the arms and legs of eight healthy adults and the difference in arm movements was measured. The SGCI values with abnormal walking patterns were compared with the SGCI values obtained during normal walking. The SCGI can be automatically and continuously measured with the gyro sensor of the smartwatch and can be used as an indirect indicator of human walking conditions.

## 1. Introduction

Walking is one of the most fundamental and primary exercises for human beings [1]. In order to determine abnormalities of walking patterns that are both directly and indirectly affected by health conditions, normal gait is often used as an indicator [2]. In other words, the occurrence of abnormal gait is a direct signal of abnormalities present in the human body. Typically, patients with Parkinson’s disease exhibit abnormal gait due to central neurological disorders. Detections of gait abnormality take a dominant role in predicting the progression of Parkinson’s disease and other neurological disorders. Recently, three-dimensional analysis of these gait changes has been conducted in various ways, and researchers have been able to digitalize gait patterns [3, 4, 5]. Nevertheless, limitations still exist. Gait analysis requires extremely expensive equipment, including multiple sensors and cameras, and cumbersome process, and thus, have constraints in measuring the long range of gait [6].

Recently, however, smart watches equipped with various heart rate sensors and physical sensors have been widely accessible to the public, improving the use of their highly developed health management functions. Furthermore, numerous studies related to the usage of smart watches in fitness have been conducted, such as measuring the time and rigor of exercises through analysis of gyro sensors and three-axis acceleration sensors. One function that this research paper will mainly focus on is the measurement of the pattern of arm swing motion, which can be depicted by the continuous graph (Figure 1).

**Figure 1.**
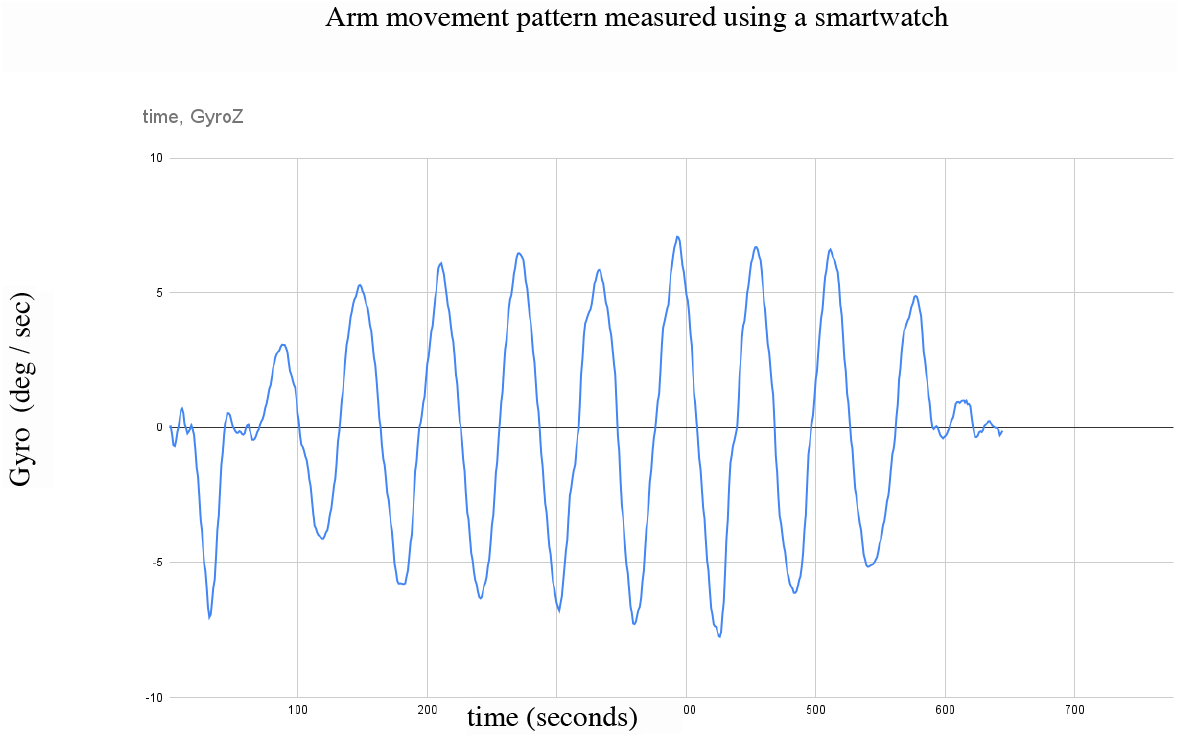
Arm movement pattern measured using a smartwatch. – This graph depicts periodic and relatively constant movement of the sinusoidal wave motion.

**Figure 2.**
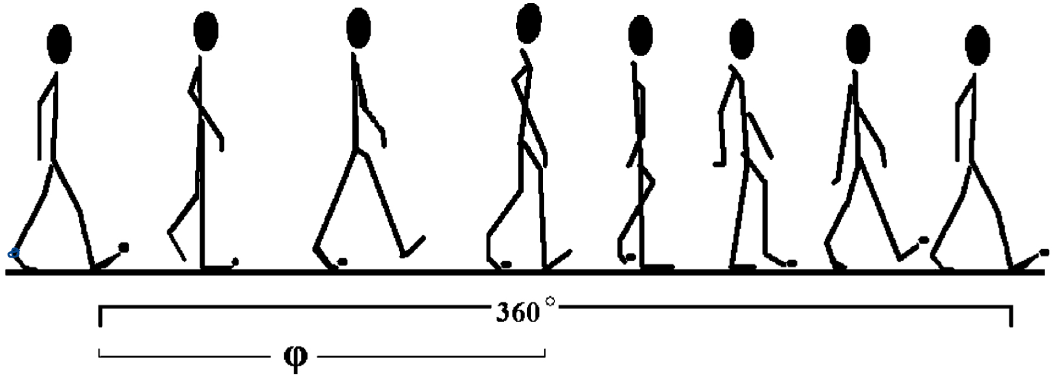
Depiction of one complete gait cycle (360 degrees) where ϕ quantifies the phase between each strike.

According to a recent study [7, 8], arm movement and walking are interrelated, meaning changes in arm swinging motions can indicate changes in walking patterns. Referencing this study, this research paper aims to analyze the arm movement patterns using acceleration and gyro sensors of smart watches to find out whether abnormalities exist in walking pattern. Furthermore, this research paper examines whether it is possible to replace the conventional analytical method with the smartwatch sensor data which quantifies the kinetic movement of the arm for detecting the walking abnormality.

## 2. Materials and Methods

### 1) Measuring smart watch gait coordination index (SGCI)

The phase coordination index (PCI) proposed by Jeffrey M. Hausdorff in 2007 [9] is a measure to examine variations of the upper limb movement by comparing the swing times of one leg to the other. PCI assumes that the walking movements of a healthy person is symmetric, meaning the phase between each strike converges to 180 degrees. Therefore, the complete cycle where one heel strike returns to that exact position on the next step is defined as 360 degrees, and the the phase of each strike is calculated by using the ratio of normalized step time and stride time :

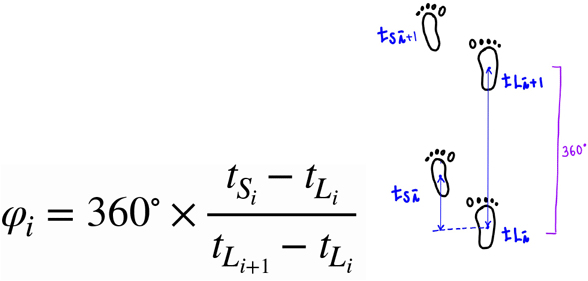

Where *t_L_i__, t_S_i__*: time of the *i*-th time the heel of the foot meets the floor.

PCI defines ϕ as the following formula, where 360 degrees — one complete cycle— is multiplied with the ratio of the time of the short distance and the long distance between heel strikes in a cycle. A person with normal walking pattern will have the value of ϕ closer to 180 degrees.

In addition, the sum of the Mean Absolute Difference (ϕ_ABS), which is the mean of absolute differences of all ϕ from the standard degree of 180, and the Coefficient of Variation (ϕ_CV) is defined as a phase coordination index (PCI) which is used as an measurement of walking motion.

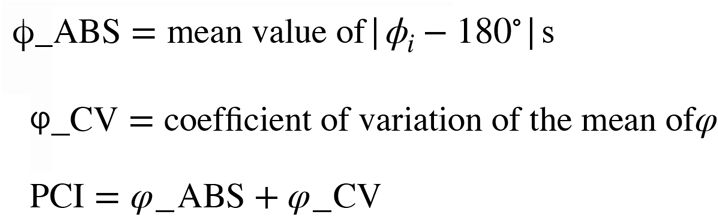

The Smart Watch Gait Coordination Index (SGCI) functions by applying PCI to the arm movement. When a person walks, both the legs and arms conduct pendulum movements. According to previous studies, it has been proven that the pendulum movement of the leg is highly correlated with the pendulum movement of the arm. When the pendulum motion of the arm is measured using a smartwatch gyro sensor, a sinusoidal wave similar to that of walking can be obtained, where one gait cycle correlates to one cycle of the arm movement’s sinusoidal wave. Therefore, the gait phase θ is defined by replacing the strike point of one foot with the maximum point of the arm movement’s sinusoidal wave and the strike point of the opposite foot with the minimum point of the arm movement’s sinusoidal wave.

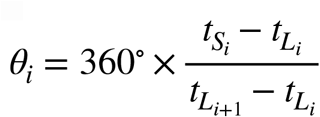

where *t_L_i__, t_S_i__*: time of the *z*-th strike point of the sine-graph with the long swing times (*L_i_*) and short swing times (*L_s_*)

SGCI, a measure of the extent of abnormality in gait motion, is defined as the sum of Mean Absolute Difference(θ_ABS) and Coefficient of Variation (θ_CV) in gait phase θ.

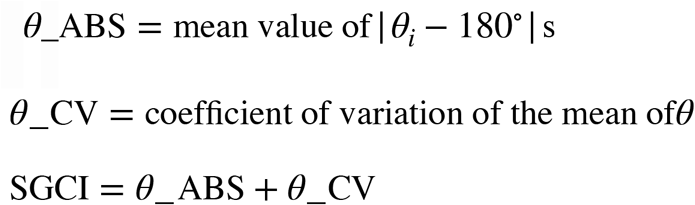

### 2) Experiments to measure arm and leg motion using smart watch

During research, we tried to prove the scientific basis of SGCI by conducting mainly two experiments: one, aiming to prove that the exercise of arms and legs are coupled when walking and another, aiming to prove whether the SGCI measured with the smartwatch can detect gait abnormality.

A gait analysis was conducted on eight healthy men whose ages were between 22-27. First, the test subjects wore a smartwatch on their wrist and ankle while walking normally, and the movements of their legs and arms were analyzed by using the data recorded on the acceleration and gyro sensors of the smartwatch. The subjects walked over eight steps on a 10 meter passage, and the walking data of the entire section was recorded. Walking data with amplitude within 10% of the maximum amplitude which continued for at least three cycles were used for actual analysis.

In addition, in order to represent abnormal walking, elements that cause walking abnormality were added to subjects normal walking patterns. More specifically, these subjects imitated people with walking abnormalities by carrying a 3kg sandbag on one ankle. Afterwards, we analyzed and compared the patterns of arm movements of walking disorders and that of normal walking.

First, we analyzed the correlation between PCI (when subjects wore smart watches on their ankle) and SGCI (when subjects wore smart watches on their wrist) during the normal walking state. Second, we obtained and analyzed the value of SGCI during abnormal walking state.

## 4. Results

In the first experiment, the measured PCI values in normal walking averaged 9.002 ± 3.872 and the 95% confidence interval for the median using the Signed Test was (2.066, 7.333). The SGCI averaged 9.847 ± 6.115, and the 95% confidence interval for the median using the Signed Test was (2.331, 9.483). The gyro sensor graph measuring the movement of the subjects’ arms and legs during normal walking state is as follows (Figure 3-1, 3-2). Both movements show stable sinusoidal waves. In fact, as a result of performing the independent t-test of both exercises’ phases measured by the strike point using the maximum and minimum values, it was confirmed that the two exercises were not statistically different as it yielded a p-value of 0.079 under the significance level *α* = 0.05.

**Figure 3-1 :**
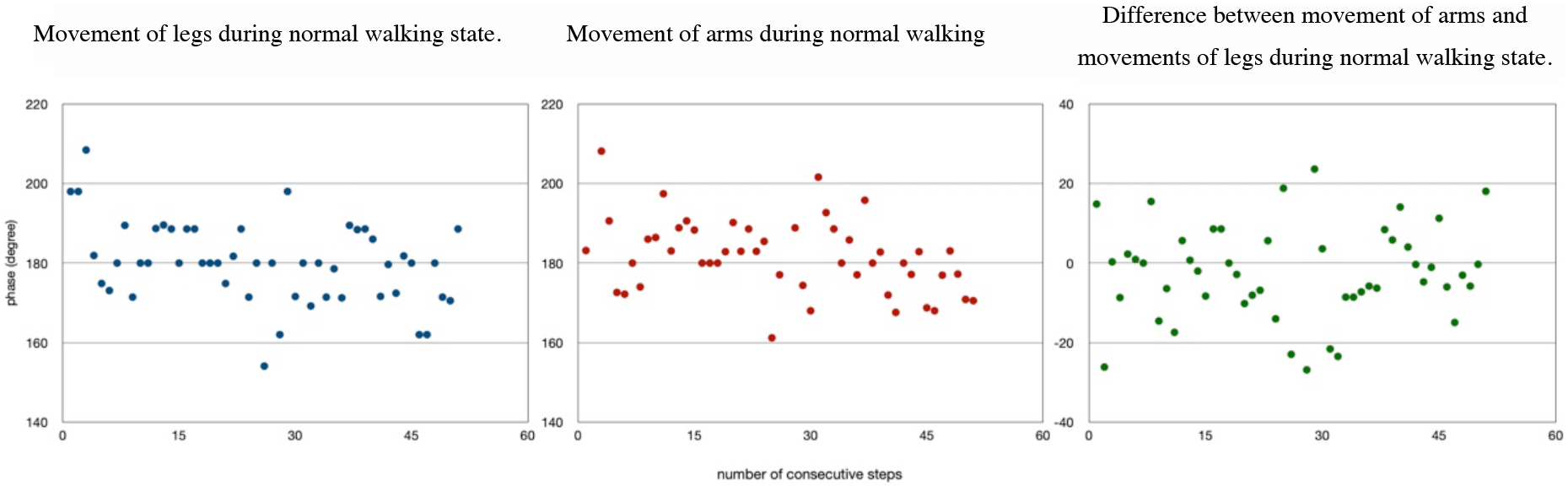
The scatter plots of phase (degrees) for consecutive steps

**Figure 3-2 :**
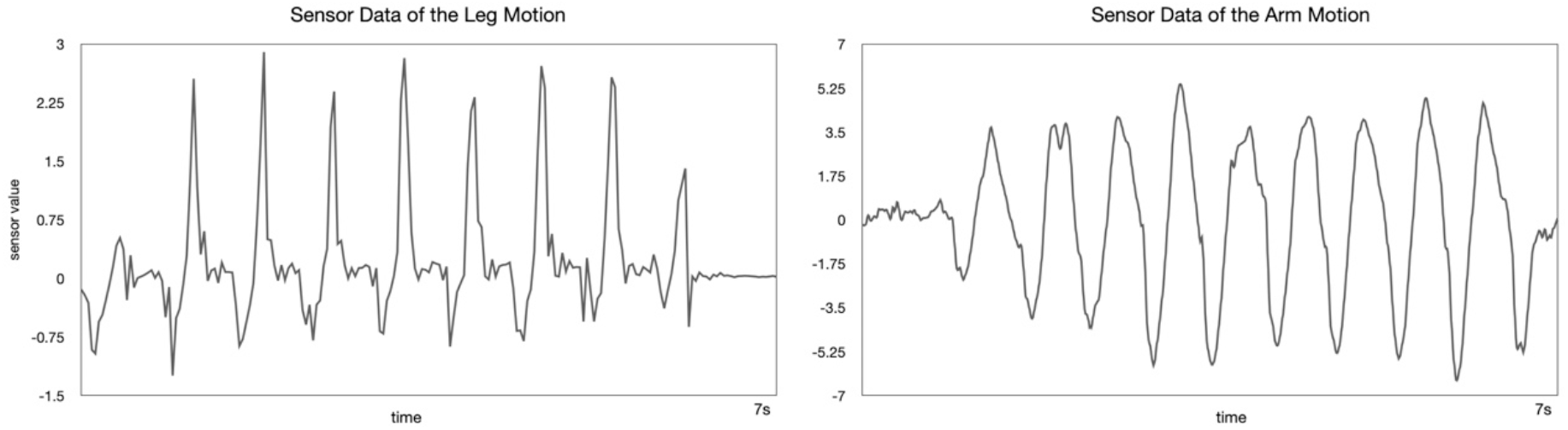
Gyro-sensor graph of leg movement (left) and gyro-sensor graph of leg movement arm (right) which show similar sinusoidal wave patterns during normal gait

The SGCI measured after applying the 3 kg weight impairment on one side leg was 22.167 ± 4.705. An asymmetric appearance of the SGCI graph, where the points are concentrated on the left, indicates that the gait phase is biased towards one side (Figure 4). Furthermore, the value of the SGCI with gait disorder conditions is statistically different from the SGCI measured without the gait disorder conditions.

**Figure 4.**
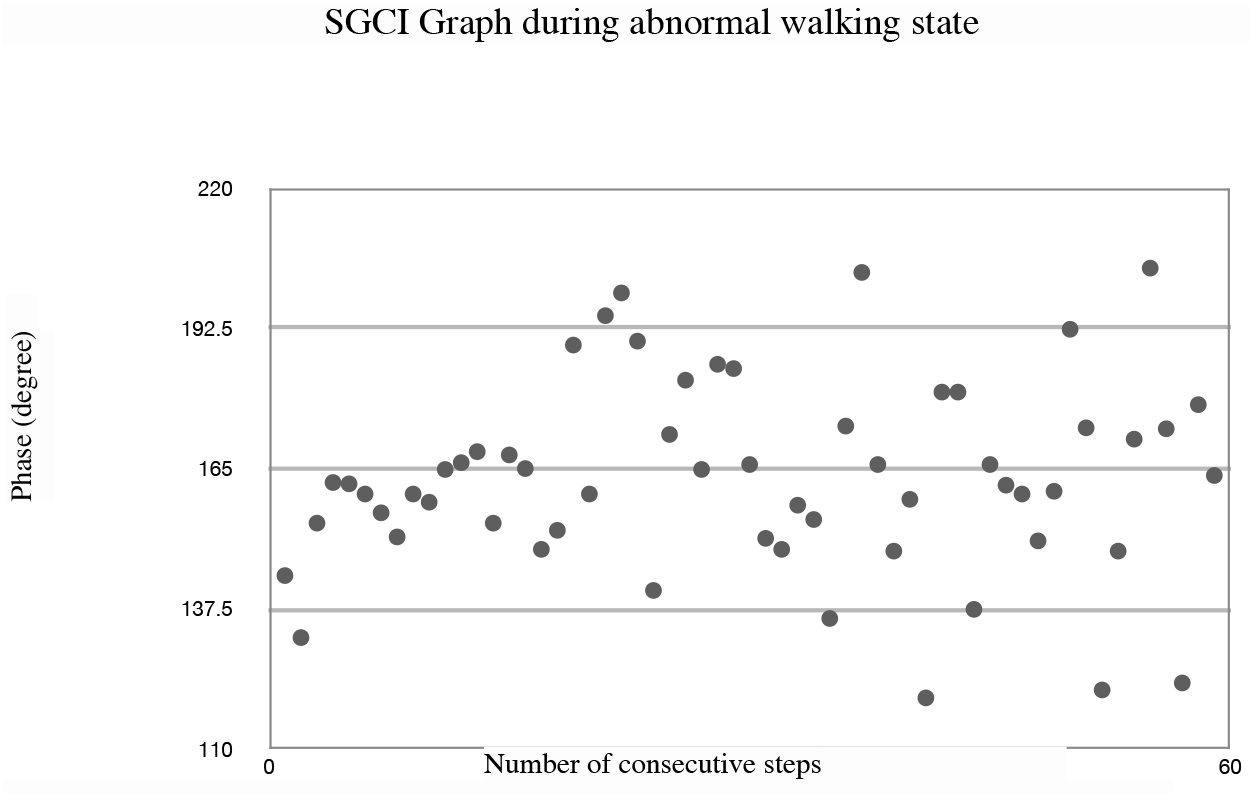
Asymmetric appearance of SCGI during disability induced gait.

## 5. Discussion

One of the simplest methods of measuring gait patterns is the method proposed by M. Jeffrey M. Hausdorff et al., which quantifies the bilateral coordination of gait using the difference between the time when the foot touches the ground and when the foot is off the ground. To summarize, from the moment when a the foot leaves the ground to the moment when it touches the ground again is defined as 2π radian, and the time ratio of the exact moment a person’s foot leaves the ground and when it touches the ground again is defined as ϕ. Using this method, the sum of Absolute Difference and Coefficient of Variation for the standard 180 degrees of one leg is defined as the Phase Coordination Index which is used as a measure of abnormality in walking motion. Currently, this measurement is applied in numerous fields for evaluating walking pattern abnormalities such as the Parkinson’s disease. However, the initially proposed PCI-calculated gait measurement method is cumbersome and quite costly. Therefore, a simple, approachable, and inexpensive method of analyzing walking is needed. As an alternative, this research paper proposes measuring walking phases using walking impairment measurement, SGCI in gyro sensors of smartwatches that are easily accessible to the public.

The SmartWatch Gait Coordination Index (SGCI) proposed in this study is an automatically measured method using smartwatch gyro sensors without the need for measurements that must be carried out by a human himself. The time used in the measurement is not when the heel touches the ground, but when the wrist reaches its highest point when walking. When a person walks using two legs, the arm exercises simultaneously according to the movement of the leg as it performs a swing motion. At this point, the wrist performs a pendulum movement with the longest radius, reaches its peak at about the same time as the heel tread, and again the pendulum swings at about the same period as the foot exercise cycle. As a result of this study, when measuring the gyro sensor of the smartwatch during walking exercise, it was found that the wrist movement was precisely reflected in the sensor, and the time when the highest point of the wrist and the maximum point of the sinusoidal wave was almost identical. Therefore, the SGCI can indirectly measure gait motion in the same way as Hausdorff’s proposed Coordination by setting the peak of the wave as the time measurement in the smartwatch gyro sensor.

It has been verified in many medical papers that the movement of the two feet and the movement of the arms during bipedalism have great connectivity with each other. When walking, the movement of the upper body, especially the movement of the arm, is directly influenced by the rotational force caused by the movement of the lower body, meaning the upper body rotates simultaneously while coordinating the movements of the legs. In particular, the rotational movement of the upper body proceeds in the opposite direction to the rotational movement of the lower body, which plays a major role in maintaining the balance of the body when walking. In a study using the actual pendulum model, it was shown that there was a high correlation between arm movement and leg movement when walking.

As a result of the experiment conducted in this study, SGCI is able to measure gait disability relatively similar to PCI, which is expected to be highly utilized in today’s world. It is possible to indirectly detect changes in walking by wearing a smartwatch in our daily life without having to visit a specific facility to measure gait disorders. Abnormal changes in SGCI may be able to selectively warn that kinematic dysfunction is present in gait. In addition, a program can be developed by applying this concept, and by installing it in a healthcare application, it will be possible to easily measure the presence and even predict the progress of gait impairment in daily life. This will reduce the inconvenience such as patients who suffer from neurological diseases or the Parkinson’s disease going to the hospital to measure the stages of the disease. Instead, SGCI can automatically determine whether an abnormality is present and correctly indicate sudden deterioration of the disease or unrecognized neurological abnormalities.

In conclusion, the SCGI can be automatically and continuously measured by using the gyro sensor of the smartwatch, and the measured SCGI can be used as an indirect indicator of the walking condition.

## References

[1] M. S. Hwang, S. G. Choi, I. K. Lee. Influence of Movement Education on Motor Development of Visually Handicapped Children. Journal of Adapted Physical Education, Vol.1, pp. 93–104, 1993.

[2] Shan DE, Lee SJ, Chao LY, Yeh SI. Gait analysis in advanced Parkinson’s disease--effect of levodopa and tolcapone. Can J Neurol Sci 2001;28:70–75.

[3] Houng S, Koh SB, Cho SC, Yoon JS, Lee SH, Park KW, et al. Analysis of dyanamics of gait in Parkinson’s disease with 3-dimensional gait analysis system. J Korean Neurol Assoc 2005;23:635–641.

[4] Nanhoe-Mahabier W, et al. Walking patterns in Parkinson’s disease with and without freezing of gait. Neuroscience. 2011;182:217–224. doi: 10.1016/j.neuroscience.2011.02.061.

[5] Nieuwboer A, Chavret F, Willems A-M, Desloovere K. Does freezing in Parkinson’s disease change limb coordination? J. Neurol. 2007;254:1268. doi: 10.1007/s00415-006-0514-3.

[6] The assessment of gait disorders in patients with Parkinson’s disease using the three-dimensional motion analysis system Vicon[.](https://pubmed.ncbi.nlm.nih.gov/17530574/)

[7] R. C. Wagenaar and R. E. van Emmerik, “Resonant frequencies of arms and legs identify different walking patterns,” J. of Biomechanics, Vol.33, No.7, pp.853–861, 2000.

[8] M. Kubo, R. C. Wagenaar, E. Saltzman, and KG. Holt, “Biomechanical mechanism for transitions in phase and frequency of arm and leg during walking,” Biological Cybernetics, Vol.91, No.2, pp.91–98, 2004.

[9] Plotnik M, Giladi N, Hausdorff JM. A new measure for quantifying the bilateral coordination of human gait: effects of aging and Parkinson’s disease. *Exp Brain Res*. 2007;181(4):561–570. doi:10.1007/s00221-007-0955-7

